# Structure-based redesign of docking domain interactions modulates the product spectrum of a rhabdopeptide-synthesizing NRPS

**DOI:** 10.1101/349555

**Authors:** Carolin Hacker, Xiaofeng Cai, Carsten Kegler, Lei Zhao, A. Katharina Weickhmann, Helge B. Bode, Jens Wöhnert

## Abstract

Several peptides in clinical use are derived from non-ribosomal peptide synthetases (NRPS). In these systems multiple NRPS subunits interact with each other in a specific linear order mediated by docking domains (DDs) to synthesize well-defined peptide products. In contrast to these classical NRPSs, the subunits of rhabdopeptide/xenortide producing NRPSs can act iteratively and in different order resulting in libraries of peptide products. In order to define the structural and thermodynamic basis for their unusual interaction patterns, we determined the structures of all N-terminal DDs (^N^DDs) as well as of an ^N^DD-^C^DD complex and characterized all putative DD interactions thermodynamically for one such system. Key amino acid residues for DD interactions were identified that upon their exchange not only changed the DD affinity but also resulted in rationally predictable changes in peptide production. A simple set of ‘recognition rules’ for DD interactions was identified that also operates in other megasynthase complexes.

## Introduction

Nonribosomal peptides are a large family of structurally diverse and pharmacologically useful natural products with broad biological activities. Prominent examples are the antibiotic daptomycin^1^ or the immunosuppressant cyclosporine A.^2^ They are assembled by multifunctional enzyme complexes called non-ribosomal peptide synthetases (NRPSs) that are organized in a modular fashion where one module activates, modifies and connects one specific amino acid with an amino acid processed by the next module. In classical NRPSs, different subunits selectively interact with each other in order to follow the collinearity rule and give rise to the synthesis of defined peptides. NRPS interactions are mediated by specialized N- and C-terminal docking domains (DDs). Stachelhaus and coworkers have demonstrated that a matching pair of DDs or COM (communication-mediating) domains at the N-terminus of peptidyl-donating NRPS (^N^DD) and the C-terminus of accepting NRPS (^C^DD) play a decisive role in defining the order of interactions between subunits *in vitro* and *in vivo* by swapping COM domains.^3^ They also constructed a “universal COM system” *in vitro* by comparison of COM domains in tyrocidin A and surfactin-like NRPSs, which led to enzyme crosstalk between different biosynthetic systems that promoted the combinatorial biosynthesis of different peptides.^4,5^ However, the structures of these DDs have not been elucidated so far but interactions between DDs have been mapped based on photo crosslinking experiments.^6^ More information on DDs is available for polyketide synthase (PKS) or PKS/NRPS hybrid systems. While for PKS three different types of DD pairs have been described structurally so far, two structures for ^N^DDs without a bound ^C^DD have been solved for NRPS/PKS hybrid systems.^7–12^

Classic NRPSs are multimodular - each protein subunit normally consists of the processing modules for multiple amino acids arranged in a linear fashion. Furthermore, the different protein subunits interact with each other in a strictly defined linear order yielding a single peptide product with a sequence that faithfully reproduces the linear order of the processing modules along the protein chains. A novel class of monomodular NRPSs involved in the production of libraries of rhabdopeptide/xenortide peptides (RXPs) has been recently described from entomopathogenic bacteria of the genera *Xenorhabdus* and *Photorhabdus*.^13^ The RXP product spectrum in a single strain differs mainly in peptide length and sequence, which requires an iterative use of certain NRPS modules and is dependent on protein stoichiometry between NRPS modules acting in elongation and termination (Supplementary Figure 1). This suggests that the subunits of these NRPSs interact with each other not in a well-defined linear order but stochastically as exemplified in the three protein model system Kj12ABC (Fig. 1). However, the sequences of the RXP products are biased suggesting that some subunits are used preferentially. These preferences might reflect differences in the interactions between subunits mediated by the DDs. The structural and thermodynamic basis for the differential docking domain interactions in these unusual NRPS systems is not clear. Therefore, we wanted to characterize docking domain interactions in these systems in detail and compare them to docking domain interactions from classical NRPS or other NRPS/PKS systems. We have solved the structures of all ^N^DDs in the three protein NRPS system Kj12ABC from *Xenorhabdus stockiae* KJ12.1.^13^ We have also characterized the thermodynamic basis for the interaction of these ^N^DDs with the two ^C^DDs present in this system and have furthermore solved the structure of one ^N^DD-^C^DD complex. The structural information for the ^N^DD/^C^DD interaction allowed us to derive a set of simple recognition rules for this type of DD interactions as well as the targeted reprogramming of selected DDs via rationally designed amino acid exchanges leading to differences in peptides produced. The type of ^N^DD/^C^DD interactions observed for this NRPS system is apparently widespread in other megasynthase systems.

**Figure 1.**
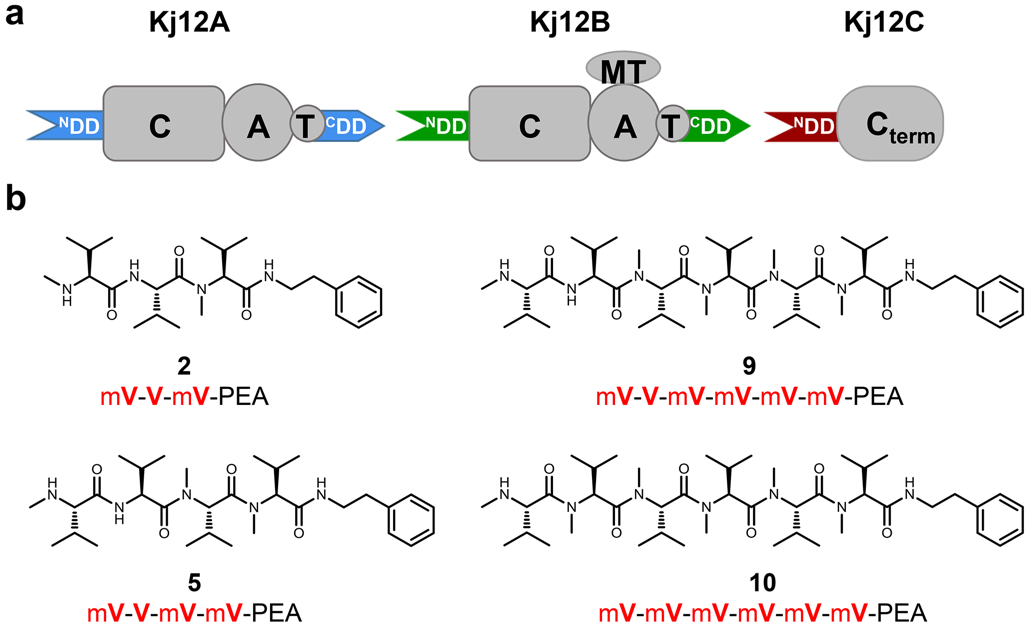
RXP NRPSs (Kj12ABC) from Xenorhabdus KJ12.1 and selected RXPs found in this strain. **a,** Overview of the domain organization of Kj12ABC (C, condensation; A, adenylation; MT, methyltransferase; T, thiolation; C_term_, terminal condensation domain; ^N^DD, N-terminal docking and ^C^DD, C-terminal docking domain). **b,** Structures of selected RXPs derived from Kj12ABC system, showing the differences in size and methylation pattern. The simplified structure nomenclature used within other figures is also shown (V, Val and mV, ***N***-methylated Val; PEA, phenylethylamine).

## Results

### Structure determination of N-terminal docking domains in monomodular NRPS of the RXP-type

Structural and thermodynamic information about docking domain interactions in monomodular NRPS systems that use their subunits in a nonlinear fashion is not available yet. The RXP NRPS system from *X. stockiae* strain KJ12.1 as a model system^13^ consists of the three proteins Kj12ABC (Fig. 1a). Bioinformatic analysis of the docking domains in this and related RXP NRPSs suggested all three proteins contained an N-terminal docking domain whereas a distinct C-terminal docking domain was found in Kj12A and Kj12B but not in the termination module Kj12C which catalyzes the reaction of the peptide chain with a terminal amine (Supplementary Fig. 2). The three ^N^DDs are ~ 65 amino acids long with more than 70% sequence identity among them (Fig. 2a and Supplementary Fig. 2a) and a predicted mixed α/β-secondary structure. Furthermore, all three ^N^DDs showed low sequence homology (< 25 % identity) to a structurally characterized ^N^DD from module TubC of the tubulysin synthesizing PKS (TubC-^N^DD) from *Angiococcus disciformis*^11^ that was shown to be a homodimer as well as to a monomeric ^N^DD of module B of the epothilone-synthesizing NRPS-PKS system (EpoB ^N^DD, 24.6 % sequence identity) crystallized in its native context as a covalent fusion with the cyclization domain of EpoB.^12^ Since the two ^C^DDs of the modules Kj12A and Kj12B were predicted to be rather short (~ 20 amino acids) and to be unstructured (Supplementary Fig. 2) we initially determined the structures of all three ^N^DDs. According to gel filtration in combination with SEC-MALS all three ^N^DDs were monomeric in solution in contrast to what was observed for the dimeric TubC ^N^DD docking domain (Supplementary Fig. 3). Solution state NMR using BEST-TROSY-based pulse sequences and non-uniform sampling rapidly yielded complete NMR resonance assignments for all three ^N^DD of Kj12ABC. The backbone chemical shift derived secondary structures for all three ^N^DDs revealed the presence of three α-helices and two β-strands in the order α1-β1-β2-α2-α3 (Fig. 2a). The location of the secondary structure elements along the sequence is very similar for the three ^N^DDs as well as to the dimeric TubC-^N^DD and the monomeric EpoB-^N^DD.^11,12^ The NMR solution structures of the three ^N^DD of the Kj12ABC were solved at very high resolution (backbone RMSD of 0.1-0.2 Å for ordered residues). A complete list of structural statistics can be found in (Supplementary Table 1) according to the recommendations of the NMR-VTF.^14^ The solution structure ensemble of the 19 lowest energy structures calculated with CYANA and an energy minimized representative mean structure for Kj12C-^N^DD is shown in Fig. 2b. A comparison of the structural ensembles and mean structures for all three Kj12-^N^DD’s are shown in Supplementary Fig. 4 In all three ^N^DDs β1 and β2 form an antiparallel β-hairpin. Helices α1 and α2 are packed against each other in an antiparallel fashion. They also pack together against one side of the β-hairpin. Helix α3 is separated only by a very short loop (aa 45) from α2 and a sharp kink is introduced in the protein backbone. Thus a3 crosses the β-hairpin in a 90° angle. Despite the very similar 3D structures (RMSDs range from 0.8 to 0.9) of the three ^N^DDs (Fig. 2c), their electrostatic surface potentials differ significantly, which could have an effect on their binding affinities for the C-terminal docking domains (Fig. 2d and Supplementary Fig. 4). A significant charge difference is found between Kj12B-^N^DD and the other two ^N^DD’s on the solvent-exposed side of strand β2 (aa 24-28) where for instance E28 in Kj12A-^N^DD and Kj12C-^N^DD is replaced by a lysine in Kj12B-^N^DD (Supplementary Fig. 4d). The topology of the three RXP-^N^DDs is already known from the TubC-^N^DD structure of *A. disciformis* (Fig. 2e),^11^ but the relative positioning of the secondary structure elements is different between the dimeric TubC-^N^DD and the RXP NRPS-^N^DDs (Cα RMSD of 4.7 Å). In contrast, the Cα RMSD is only 1.3 Å (Supplementary Fig. 4) between Kj12C-^N^DD and the monomeric EpoB-^N^DD.

**Figure 2.**
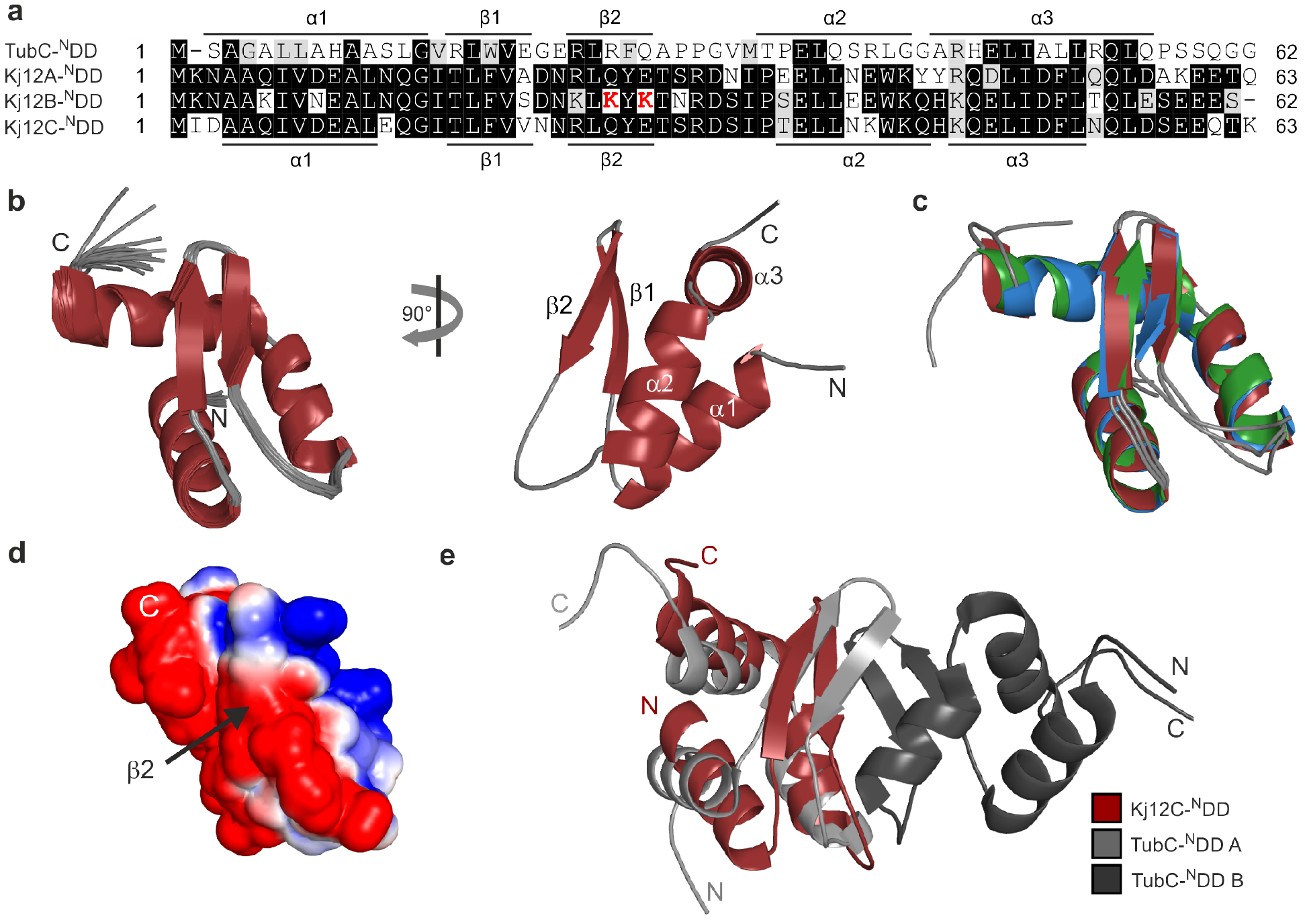
N-terminal docking domains have the same three-dimensional structure. **a,** Structure based sequence alignment of the TubC-^N^DD and the N-terminal docking domains of Kj12ABC. Identical residues are highlighted with *dark gray boxes* and residues with similar chemical properties are shown in light gray boxes. Key residues for DD interactions are shown in red. The secondary structure based on structural information for TubC-^N^DD and Kj12C-^N^DD are indicated above and below the sequence. **b,** Solution structure bundle of the 19 lowest energy conformers and the regularized mean structure for Kj12C-^N^DD. **c,** Overlay of the cartoon representations of the energy minimized mean structures of Kj12A-^N^DD (blue), Kj12B-^N^DD (green) and Kj12C-^N^DD (red). **d,** Electrostatic surface potentials of Kj12C-^N^DD mapped on the solvent accessible surfaces in the same orientation as in b (left) with negatively charged surface areas colored in red, positively charged areas coloured in blue and white areas corresponding to hydrophobic surfaces. **e,** Overlay of the energy minimized mean structure of Kj12C-^N^DD with one monomer of the TubC-^N^DD dimer.

### Interactions with the ^C^DDs

To further investigate the docking domain interaction NMR titration experiments were carried out. Therefore, all three ^15^N labeled ^N^DD proteins were titrated with both unlabelled (^14^N) ^C^DD peptides (Fig. 3a). In Fig. 3b the titration of Kj12C-^N^DD with Kj12B-^C^DD as followed in ^1^H,^15^N-HSQC-experiments is exemplarily shown. The NMR-data for all other titrations are shown in Supplementary Fig. 5. In all six titration experiments gradual chemical shift changes and/or peak broadening during the stepwise addition of the ^C^DD peptides are observed. This suggests that all three ^N^DDs bind to both the Kj12A- and the Kj12B-^C^DD in agreement with the observed product spectrum of this NRPS and form ^N^DD/^C^DD complexes in the fast to intermediate exchange regime on the NMR time scale. The chemical shift changes during the titrations were quantified and mapped on the structures of the three ^N^DDs in order to identify the ^C^DD binding sites. A histogram of chemical shift changes vs. sequence for the titration of the Kj12C-^N^DD with the Kj12B-^C^DD (Fig. 3c) and a mapping of these shift changes on the cartoon representation of the Kj12C-^N^DD structure is shown in Fig. 3d (also see Supplementary Figs. 6 and 7 for the data of all six titrations). Importantly, in all six titration experiments, the largest chemical shift changes are always observed for amino acids in the β hairpin and in particular strand β2 as well as helix α2 of the three ^N^DDs. This suggests that both the Kj12A-and Kj12B-^C^DD bind to all three ^N^DDs at the same site.

The pairwise interactions between the three ^N^DDs and the two ^C^DDs were further characterized thermodynamically using isothermal titration (ITC) experiments (Supplementary Fig. 8 and Supplementary Table 2). The obtained K_d_ values for all interactions (Fig. 3e) are in good agreement with the fast to intermediate exchange observed in the NMR titrations. The highest affinity interactions with 8 ± 4 μM and 8 ± 6 μM were observed between the Kj12B-^C^DD and the ^N^DDs of Kj12A and Kj12C, respectively. The Kj12B-^N^DD binds Kj12B-^C^DD with an ~8-fold higher Kd of 62 ± 8 μM. Kj12A-^C^DD (Fig. 3e) binds only weakly to Kj12C-^N^DD (~100 μM) whereas its interaction with the other two ^N^DDs is too weak to be reliably quantified (Supplementary Fig. 8) The reported affinities for docking domain pairs in other megasynthases were found to be in a similar range as those observed here.^9,11^

**Figure 3.**
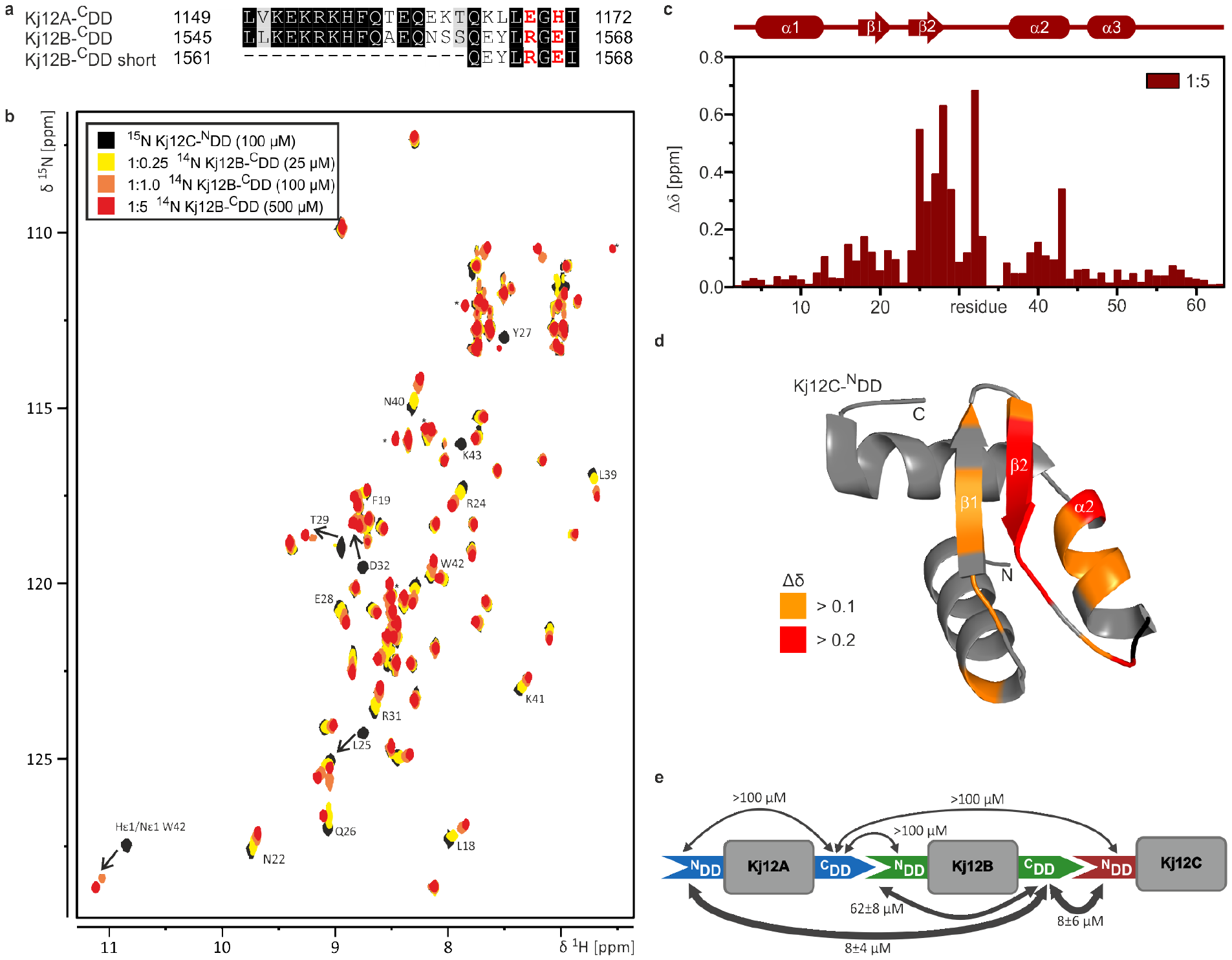
Docking domain interaction. **a,** Sequence alignment of Kj12A-^C^DD and Kj12B-^C^DD used in this study. Identical residues are highlighted with dark gray boxes and residues with similar chemical properties are shown in light gray boxes. Key residues for DD interactions are shown in red. **b,** Overlay of the ^15^N-HSQC spectrum of 100 ^15^N labeled Kj12C-^N^DD in the absence (black) and presence of increasing amounts of unlabeled Kj12B-^C^DD. The molar ratios of the two docking domains are 1:0.25 (yellow), 1:1 (orange), 1:2.5 (dark orange), 1:5 (red). **c,** Histogram of the chemical shift changes vs. sequence of Kj12C-^N^DD upon addition of a 5-fold molar excess of Kj12B-^C^DD in its unlabeled form with the secondary structure depicted above. **d,** Chemical shift changes of Kj12C-^N^DD upon addition of unlabeled Kj12B-^C^DD mapped onto the structure of Kj12C-^N^DD. **e,** Kd values for all DD interactions in the Kj12ABC NRPS as measured with ITC.

### Solution Structure of a Docking Domain Complex

Structural information about the interaction of the type of ^N^DD found in the Kj12 RXP NRPS cluster so far is limited to ^C^DD peptide titration experiments for the dimeric TubC-^N^DD where the interaction surface on the dimeric ^N^DD was identified but no complex structure was obtained.^11^ Furthermore, our NMR titration experiments identify a ^C^DD binding site on the monomeric ^N^DDs that is part of the dimer interface in the TubC-^N^DD. Thus, the structural basis for the ^N^DD/^C^DD interactions in the Kj12 RXP NRPS is apparently different from what was observed for TubC. For this reasons and in order to understand the underlying reasons for the widely different ^N^DD/^C^DD affinities, we exemplarily solved the structure for one ^N^DD/^C^DD complex from the RXP NRPS. We chose Kj12C-^N^DD and Kj12B-^C^DD to solve a docking domain complex structure since we observed a comparatively high affinity for this interaction. This complex is however, still in fast to intermediate exchange on the NMR time scale and therefore the collection of a large enough number of intermolecular NOEs is difficult. To overcome this problem, we designed a covalently linked ^N^DD-^C^DD complex with flexible glycine-serine linkers of different lengths to increase the local ^C^DD concentration at the ^N^DD. To verify that our artificially linked DD pair interacts in *cis* with the same binding mode as the isolated domains in *trans* we compared the titration endpoint ^1^H,^15^N-HSQC spectrum of the separate domains with the ^1^H,^15^N-HSQC spectrum of the linked constructs (Supplementary Figs. 9 and 10). For this purpose, we screened different constructs with different linker length (6, 9 and 12 residues, data not shown for 6 and 9 residue linker) and different domain order (^N^DD-linker-^C^DD or ^C^DD-linker ^N^DD). The construct with the longest linker (12 residues) and an ^N^DD-^C^DD arrangement (Fig. 4a) was the best mimic for the natural ^N^DD-^C^DD complex and therefore we solved the structure of this fusion protein (Fig. 4b). To our knowledge, this structure represents the first high-resolution structure of a NRPS docking domain pair. Surprisingly, the ^N^DD-^C^DD interaction involves only the last five C-terminal amino acids of the ^C^DD. These amino acids form an additionally β-strand, b3 which interacts in an antiparallel orientation with β-sheet, b2 of the ^N^DD as well as with parts of helix α2 of the ^N^DD (Fig. 4c and Supplementary Fig. 11). The additional α-helix formed by the first nine amino acids of the ^C^DD does not interact with the ^N^DD (Fig. 4b). The overlay with the structure of the isolated ^N^DD shows that the ^N^DD does not change its conformation upon binding to the ^C^DD (Supplementary Fig. 12).

A detailed view of the “intermolecular” backbone hydrogen bonding interactions between β-strands β2 and β3 from the ^N^DD and the ^C^DD, respectively is shown in Supplementary Fig. 13. Interestingly, the C-terminal end of the β-sheet of Kj12B ^C^DD (β3) is highly twisted towards helix α2 (Fig. 4c). Thereby, the side chain of the last amino acid (I1568) is buried in a hydrophobic pocket consisting of the surrounding side chains of helix α2 of the ^N^DD (Supplementary Fig. 11). The interaction of the ^C^DD with this helix explains the large chemical shift changes (Supplementary Fig. 6) observed for aa 39-43 of the ^N^DD during the titration experiments. Additionally, the side chain of the first amino acid of β3 (L1564) is located in a hydrophobic pocket build by hydrophobic sidechains from β2 and from the loop between β2 and α2 (Supplementary Fig. 11). The signals for the backbone amide groups of the loop residues are either not observable (I34) or have very low intensities in the free ^N^DD (D32, S33) indicative of conformational exchange. In the complex with the ^C^DD these signals have significantly higher intensities. Thus, loop2 of the ^N^DD is stabilized upon ^C^DD binding. Y27 of the ^N^DD packs tightly against the small side chain of G1566 in the ^C^DD. The “intermolecular” interaction between β2 of the ^N^DD and β3 of the ^C^DD is also stabilized by two salt bridges between side chains involving R24 of β2 and E 1567 of β3 as well as E28 of β2 and R1565 of β3, respectively (Fig. 4c).

**Figure 4.**
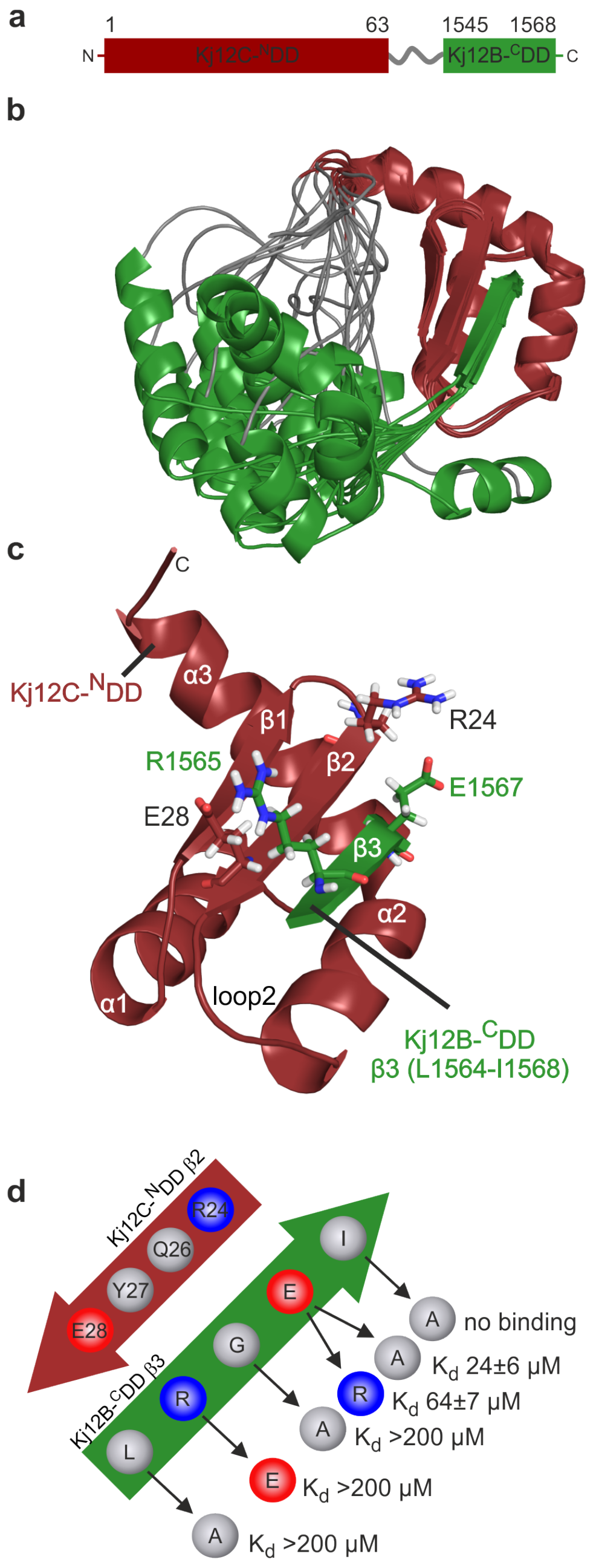
Structure of ^N^DD-^C^DD complex. **a,** Schematic representation of the N-and C-terminal docking domain linker construct used in this study with Kj12C-^N^DD in red, the 12 amino acid long Gly-Ser linker in grey and Kj12B-^C^DD (amino acids 15451568 from Kj12B corresponding to residues 75-99 in the complex construct Kj12C-^N^DD-12xGS-Kj12B-^C^DD) in green. **b,** Solution structure bundle of Kj12C-^N^DD-Kj12B ^C^DD linker construct with colour coding as in a. **c,** Detailed view of the ^N^DD-^C^DD interaction of β-sheet 2 of Kj12C-^N^DD (red) with the last 5 amino acids of Kj12B-^C^DD (green), stick representation of charged residues forming salt bridges between the two docking domains. **d,** Schematic representation of the ‘recognition rules’ for the interaction of β2 of Kj12C-^N^DD (red) and β3 of Kj12B-^C^DD (green) in the complex. Positively and negatively charged residues are shown as blue and red circles, respectively, and hydrophobic residues as white circles. Kd values were determined by ITC titration experiments with synthetic peptides of Kj12B-^C^DD_short_ carrying variations of all five residues of β3.

Taken together, ^N^DD/^C^DD complex formation is apparently dependent upon the formation of an intermolecular β-sheet stabilized by two salt-bridges and the burial of two large hydrophobic side chains. The involvement of only the five C-terminal amino acids but not the remainder of the ^C^DD in complex formation is also supported by hetNOE measurements. In agreement with the complex structure, they show low values for residues in the GS-linker and in the majority of the ^C^DD suggesting that these amino acid residues are flexible in solution. Only the C-terminal amino acids of the ^C^DD which are part of β3 have the same high hetNOE values as observed for the conformationally rigid residues of the ^N^DD (Supplementary Fig. 14). To further verify that the ^N^DD-^C^DD interaction is exclusively based on the β-sheet interaction we repeated the titration experiments with a shortened Kj12B-^C^DD peptide which only comprises the β-sheet (eight C-terminal residues from Kj12B-^C^DD, Fig. 3a). The endpoint of both NMR titration experiments (Kj12C-^N^DD with Kj12B-^C^DD and Kj12B-^C^DD_short_, respectively, in identical concentrations and ratios) were perfectly overlapping (Supplementary Fig. 15) and ITC measurement confirmed that the short ^C^DD peptide binds with the same affinity (10 ± 3 μM) as the original longer ^C^DD peptide (Supplementary Fig. 15).

In order to test the relative importance of the observed intermolecular interactions variants of the Kj12B-^C^DD_short_ peptide were synthesized (Supplementary Fig. 16) and tested. Replacement of either of the two large hydrophobic side chains L1564 and I1567 by the smaller alanine lead to a significant loss of affinity (Supplementary Table 2). Increasing the size of the side chain of G1565 which is stacking against Y27 by replacement with alanine also leads to a decrease in binding affinity. Importantly, breaking of the salt-bridges between E28 and R1565 and R24 and E1567, respectively, significantly lowers the affinities between the Kj12C-^N^DD and the ^C^DD peptide. In particular, the salt bridge between E28 and R1565 seems to contribute strongly to the binding affinity.

In this respect it is interesting to note that in the Kj12B-^N^DD E28 is replaced by lysine (Fig. 2a) explaining the lower affinity measured for its interaction with the native Kj12B-^C^DD peptide. In contrast, a peptide R1565E (Fig. 3a) that would restore salt bridge formation with K28 in Kj12B-^N^DD binds with a much higher affinity to the Kj12B-^N^DD (13 ±1 μM). Furthermore, these data rationalize why the Kj12A-^C^DD binds rather weakly to all three ^N^DDs. It contains an H at the position corresponding to E1567 in Kj12B-^C^DD and an E at the position corresponding to R1565 in Kj12B-^C^DD (Fig. 3a), thereby weakening or preventing the formation of either one of the two intermolecular salt bridges. However, these data also suggest that tuning of the intermolecular docking domain salt-bridge interactions in the context of the full-length modules could influence the interactions of the modules and thereby the product spectrum in this NRPS system.

### DD-reprogramming results in peptide elongation and diversification

In order to verify the identified key residues for protein-protein interaction in the Kj12ABC system, we systematically changed selected positions in the DDs. As a starting point, the Kj12A-^C^DD was changed to E1169R and H1171E in order to increase the affinity between Kj12A-^C^DD and Kj12C-^N^DD. Indeed, the optimized Kj12A variant was able to interact with Kj12C resulting in the formation of RXP **1**, V-PEA not detected in the native KJ12AC system (Supplementary Fig. 17). When the optimized Kj12A was combined with native Kj12BC an increase of longer RXPs was observed (Fig. 5a and Supplementary Fig.18) that was also observed for an artificial system where the amino acid specificity of Kj12B was changed from V to L allowing an easier differentiation of the activities of Kj12A and Kj12B (Supplementary Fig. 19). This might result from efficient binding between Kj12A and Kj12C so that Kj12C is not available for peptide termination via binding to Kj12B anymore since it was previously shown that the peptide length is dependent on the protein stoichiometry between Kj12B (elongation) and Kj12C (termination).^13^

A similar production of longer peptides with a chain length of up to 10 amino acids was observed in a Kj12BC system when the Kj12B-^N^DD was optimized via K28E, K26Q and K24R exchange for a higher affinity towards Kj12B-^C^DD (Fig. 5b and Supplementary Fig. 20) as it was also shown in ITC measurements (Supplementary Fig. 21). Amino acid exchange of E28A and Q26K in the ^N^DD of Kj12C designed to weaken the interaction between the Kj12B-^C^DD and the Kj12C-^N^DD and thereby to reduce termination also resulted in longer peptides of up to nine amino acids in a Kj12BC system (Supplementary Fig. 22). When additionally the optimized Kj12B-^N^DD (K28E, K26Q and K28E) and Kj12C-^N^DD (E28A and Q26K) were combined even more long-chain peptides were produced (Fig. 5c and Supplementary Fig. 22).

**Figure 5.**
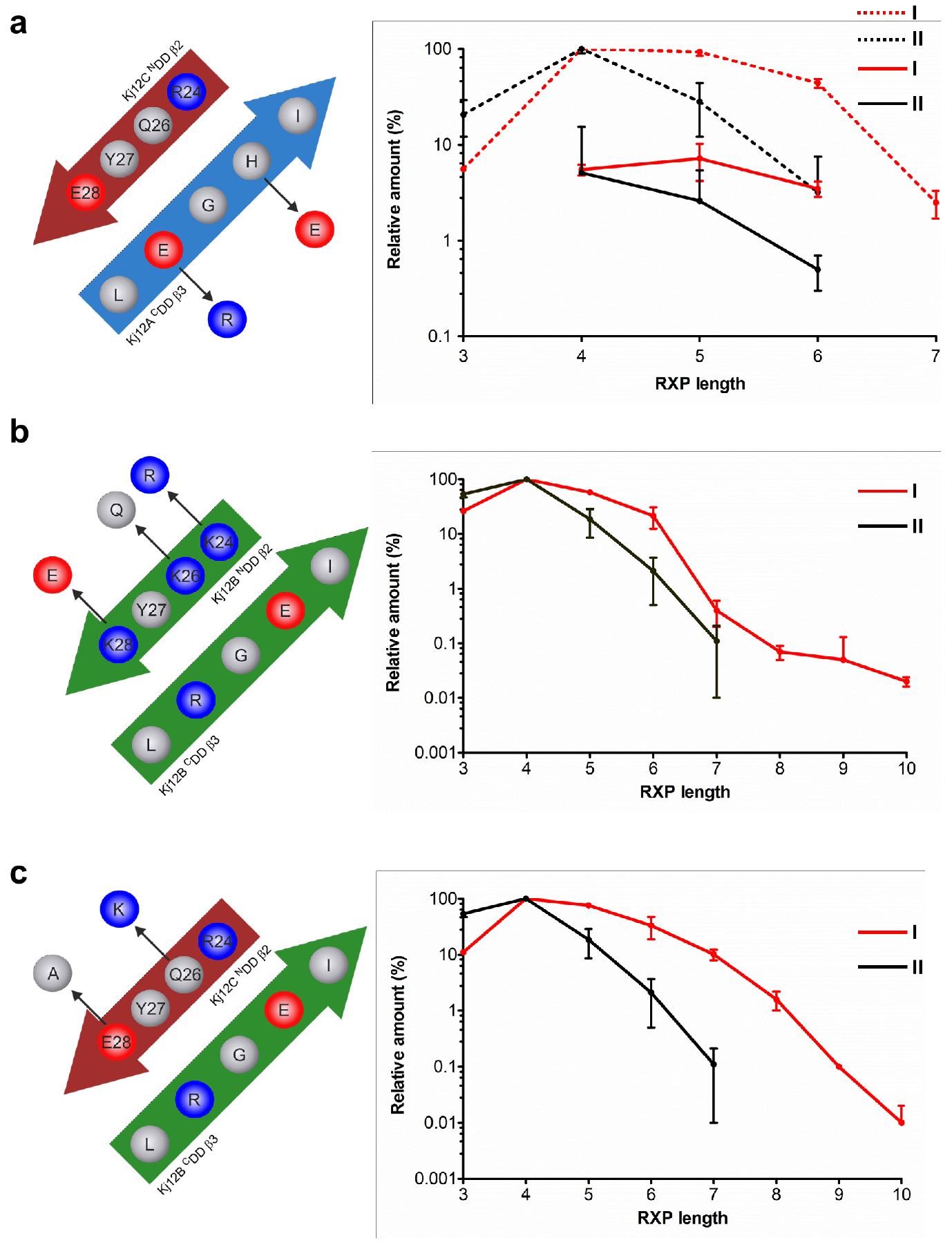
Optimization of ^C^DD or ^N^DDs in Kj12ABC system for production of longer RXPs. **a,** Kj12A-^C^DD was optimized by two amino acid exchanges, E1169R and H1171E on Kj12A-^C^DD. Co-expression of natural Kj12B with optimized Kj12C led to increased production of longer RXPs. Red lines (I) represent RXP production in the modified system, black lines (II) represent RXP production in the natural Kj12BC. Solid lines indicate fully methylated Val (mV) RXPs, dash lines indicate RXPs containing only one non-methylated Val. **b,** Optimization of Kj12B-^N^DD for better interaction with Kj12B-^C^DD via three amino acid exchanges, K26Q, K24R and K28E on Kj12B-^N^DD. Red line (I) represents RXP production in the optimized system, black line (II) represents RXP production in the natural Kj12BC. Only fully methylated RXPs were shown. **c,** in addition to amino acid exchanges in **b,** Kj12C-^N^DD was additionally modified via two amino acid exchanges, Q26K and E28A to reduce the interaction with Kj12B-^C^DD and allowing better interaction between Kj12B-^N^DD and Kj12B-^C^DD. No RXPs were detected after co-expression of natural Kj12B and modified Kj12C-^N^DD probably due to very weak affinities. Red line (I) represents RXP production in the optimized system, black line (II) represents RXP production in natural Kj12BC. Only fully methylated RXPs were shown. X axis, Numbers of amino acid residues in RXPs (RXP length). Y axis, production of the corresponding RXPs relative to the most abundant derivative set to 100%.

### The DD ‘recognition rules’ can be applied to predict DD interactions in many other megasynthase systems

In order to investigate whether the DD recognition mode of the RXP-NRPS system observed here also occurs in other systems, a BLASTP search using Kj12C-^N^DD and TubC-^N^DD as query sequence was carried out. Indeed, related DDs were identified (Supplementary Figure 2) and putative interactions between ^N^DDs and ^C^DDs dependent on salt bridges and hydrophobic interactions similar to the Kj12ABC system could be predicted (Supplementary Figure 23). However, in some examples the polarity of the salt bridges is reversed. In other cases a salt bridge is replaced by a hydrogen bond interaction or by a hydrophobic interaction. Besides highly related RXP-NRPS systems, a new family of NRPS related to the taxlllaid lipopepide-producing NRPS TxlAB^15^ has been identified in *Xenorhabdus* strains. Here the DDs connecting NRPS subunits carrying C-terminal epimerization (E) to N-terminal condensation (C) domains (Supplementary Figure 2) interact in a fashion similar to that observed in Kj12ABC.

Additionally, DD pairs with a very similar putative interaction mode were identified in *Pseudomonas, Janthinobacterium, Paenibacillus,* cyanobacteria and myxobacteria (Supplementary Figures 2 and 23). While the natural products for *Pseudomonas, Janthinobacterium* and *Paenibacillus* are not known yet, they can be predicted to be two novel peptides and a peptide/polyketide hybrid (*Paenibacillus*) as determined from an antismash analysis^16^ of their genomes. In cyanobacteria the biosynthesis machineries for the peptides micropeptin^17^ and cyanopeptolin^18^ and in myxobacteria for the peptide/polyketide hybrid melithiazol^19^ were identified.

## Discussion

Engineering of NRPS or PKS systems for the production of derivatives or even new natural products requires the efficient modification of catalytic domains^20,21^ but also the modification of protein-protein interaction of megasynthases^22^ consisting of multiple enzymes as is the case for most systems that incorporate >5 building blocks.

Here we have analyzed the DD-mediated interactions required for the production of rhabdopeptides using the RXP type of monomodular NRPS systems from *Xenorhabdus* KJ12.1 as a model system. We could show that amino acid exchanges in the respective DDs not only results in the predicted shift in protein affinity *in vitro* but also in the production of different peptides *in vivo*.

From the overlay of the TubC-^N^DD structure with Kj12C-^N^DD it is obvious that the binding sites for the ^C^DDs must be very different in the two systems. This is due to the dimeric structure of the TubC-^N^DD where the β-hairpin is part of the dimerization surface (Fig. 2e) and provides no space for the β-sheet interaction with a ^C^DD observed in Kj12ABC. However, from a brief analysis of different biosynthesis gene clusters it was obvious that several other megasynthases exist that could use ^N^DD/^C^DD interactions that are structurally similar to what we found for Kj12ABC (Supplementary Figures 2 and 23) and conform to similar recognition rules.

The understanding of megasynthase DDs is a basis for future engineering approaches of such systems since it might allow the combination of different proteins that have been individually optimized in a combinatorial approach. Also the splitting of large megasynthases too difficult to engineer might be possible using well-studied DDs. Additionally DD enginered cross-talk of megasynthase subunits from different biosynthesis pathways could even increase the chemical diversity beyond natural NPs and the DD specificity code described here allows the fast identification of specific protein-protein interactions and thus might help to elucidate biosynthesis pathways for systems that are not collinear. Moreover, from our analysis it is also obvious that additional DD types exist for NRPS and other megasynthases that require further structural and biochemical analysis allowing their future use in NRPS engineering or understanding the basic principles of these megasynthase pathways.

## Data availablity statement

All structures have been deposited in the Protein Data Bank under the ID codes 6EWS (Kj12A-^N^DD), 6EWT (Kj12B-^N^DD), 6EWU (Kj12C-^N^DD) and 6EWV (Kj12-^N^DD-Kj12B-^C^DD).

## Supporting Information

Experimental procedures, chemical synthesis of all peptides are included in the supporting information.

**Notes** The authors declare no competing financial interests.

## Acknowledgement

This work was supported by the LOEWE program (Landes-Offensive zur Entwicklung wissenschaftlich-ökonomischer Exzellenz) of the state of Hesse and was conducted within the framework of the MegaSyn Research Cluster in the labs of HBB and JW. Additionally, work in the Bode lab was supported by an ERC Starting Grant (grant agreement number 311477). All NMR experiments were carried out at the Center for Biomolecular Magnetic Resonance (BMRZ) at Goethe-University Frankfurt, which is supported by the state of Hesse. We are grateful to Dr. Frank Löhr and Dr. Christian Richter for their support in setting up BEST-TROSY-based experiments and to Dr. Elke Duchardt-Ferner for helpful discussions.

## Author contributions

C.H. and X.C. contributed equally to this work. H.B.B. and J.W. designed the experiments. C.H. performed all NMR and ITC experiments and elucidated the DD structures. Constructs for the production of DDs were performed by C.K. and X.C.. X.C. performed all *in vivo* experiments including strain construction and natural product quantification. Synthesis of ^C^DDs was performed by L.Z. A.K.W. performed ITC experiments. C.H., X.C., J.W. and H.B.B. wrote the paper with input from all authors.

